# Patient-derived cancer organoid tracking with widefield one-photon redox imaging to assess treatment response

**DOI:** 10.1101/2020.12.19.423617

**Authors:** Daniel A. Gil, Dustin Deming, Melissa C. Skala

**Affiliations:** Department of Biomedical Engineering, University of Wisconsin, Madison, WI; Morgridge Institute for Research, Madison, WI; University of Wisconsin Carbone Cancer Center, Madison, WI; Division of Hematology and Oncology, Department of Medicine, University of Wisconsin, Madison, WI; McArdle Laboratory for Cancer Research, University of Wisconsin, Madison, WI; William S. Middleton Memorial Veterans Hospital, Madison, WI

**Keywords:** autofluorescence, redox imaging, image analysis, tracking, drug screening, cancer organoid

## Abstract

**Motivation:** Accessible tools are needed for rapid, non-destructive imaging of patient-derived cancer organoid (PCO) treatment response to accelerate drug discovery and streamline treatment planning for individual patients.

**Aim:** Segment and track individual PCOs with widefield one-photon redox imaging to extract morphological and metabolic variables of treatment response.

**Approach:** Redox imaging of the endogenous fluorophores, NAD(P)H and FAD, was used to monitor the metabolic state and morphology of PCOs. Redox imaging was performed on a widefield one-photon epifluorescence microscope to evaluate drug response in two colorectal PCO lines. An automated image analysis framework was developed to track PCOs across multiple time points over 48 hours. Variables quantified for each PCO captured metabolic and morphological response to drug treatment, including the optical redox ratio and organoid area.

**Results:** The optical redox ratio (NAD(P)H/(FAD+NAD(P)H)) was independent of PCO morphology pre-tieatment. Drugs that induced cell death decreased the optical redox ratio and growth rate compared to control. Multivariate analysis of redox and morphology variables identified distinct PCO sub-populations. Single-organoid tracking improved sensitivity to drug treatment compared to pooled organoid analysis.

**Conclusion:** Widefield one-photon redox imaging can monitor metabolic and morphological changes on a single organoid-level, providing an accessible, non-destructive tool to screen drugs in patient-matched samples.

## 1 Introduction

Precision medicine aims to improve cancer treatment by matching optimal therapies for each patient, typically based on genomic mutations.^1–3^ While effective for a subset of mutations (e.g., EGFR in lung cancer, BRAF in melanoma), this approach has not been as successful in all cancers.^4^ Alternatively, drugs can be directly tested on a patient’s tumor cells, providing functional information on drug sensitivity that is complementary to genomic approaches.^4^ Patient-derived cancer organoids (PCOs) are 3D organotypic cultures grown from fresh tumor samples (e.g., surgery, biopsy). PCOs recapitulate the *in vivo* molecular, histopathological, and phenotypic features of the original patient tumor.^5–7^ PCOs also provide accurate models of patient drug sensitivity, so that multiple drugs can be screened in patient-matched samples for streamlined drug development and clinical treatment planning.^5,6,8–11^ Additionally, PCOs capture the cellular heterogeneity found in tumors, which can result in treatment failure if drug resistant cell subpopulations are present.^5,10,12,13^ Therefore, it is critical to measure drug response across multiple PCOs to capture the response of sub-populations and accurately predict patient response. However, current tools to evaluate drug response in PCOs are destructive, providing only a static measurement that does not capture the dynamic nature of drug response and prohibits the use of complementary assays. This highlights a significant need to quantify drug response in PCOs using non-destructive tools that are sensitive to cellular heterogeneity.

Cellular metabolism is altered in cancer, and most cancer therapies disrupt cellular metabolism to limit proliferation or induce apoptosis.^14^ Autofluorescence can non-destructively monitor cellular metabolism within intact samples through the endogenous, metabolic co-enzymes nicotinamide dinucleotide (NADH), nicotinamide dinucleotide phosphate (NADPH), and flavin adenine dinucleotide (FAD).^15–23^ NADH and NADPH have overlapping fluorescent properties and are jointly referred to as NAD(P)H. NAD(P)H and FAD are involved in hundreds of metabolic reactions, including glycolysis and oxidative phosphorylation. The fluorescence intensities of NAD(P)H and FAD can be combined into an “optical redox ratio” [i.e., NAD(P)H/(FAD+NAD(P)H)] that provides a per-pixel map of the oxidation-reduction state of a sample.^24–26^ The optical redox ratio is sensitive to early drug-induced changes in cell metabolism that precede changes in tumor volume, proliferation (e.g., Ki67), and cell death (e.g., cleaved caspase 3).^26–32^ However, most studies of autofluorescence in PCOs have used expensive multiphoton and/or fluorescence lifetime imaging microscopes that are not widely available.^26–32^ A recent study by our group used selective plane illumination (light sheet) microscopy (SPIM) to image the autofluorescence of multiple PCOs, but required non-standard culture dishes to accommodate the imaging geometry of the microscope.^33^ These drawbacks complicate the application of these techniques in drug discovery or the clinic. Disseminating autofluorescence screens beyond specialized laboratory use requires validation of accessible instrumentation that is compatible with standard cell culture methods.

Redox images can be collected with commonly available widefield one-photon epifluorescence microscopes, which generally include a broadband excitation source, scientific monochrome camera, and standard filter cubes (DAPI and GFP cubes were used for NAD(P)H and FAD, respectively).^15^ In this widefield configuration, redox imaging can capture the optical redox ratio of each pixel along with the morphology of each PCO, but this configuration does not resolve individual cells as in previous two-photon redox imaging studies.^26–32^ However, widefield one-photon redox imaging can rapidly image large populations of PCOs, which is important for characterizing sub-populations of PCOs with variable treatment sensitivities.^9,34^ New image analysis methods are needed for widefield one-photon redox imaging that can resolve drug response in heterogenous PCO populations. Specifically, image segmentation and tracking methods are needed to quantify the drug response of each PCO over a treatment time course. These methods enable quantitative measures of the drug response (e.g., changes in organoid morphology and optical redox ratio) of each PCO with respect to its matched pre-treatment values. Statistical analysis techniques, such as mixed-effect models, can leverage these longitudinal data to account for organoid-level heterogeneity and provide improved sensitivity to drug-induced changes in redox imaging variables.

Here, we provide the first demonstration of widefield one-photon redox imaging of PCOs. A novel image analysis framework was developed to automatically quantify organoid-level changes in redox ratio, autofluorescence intensity, and morphology over a treatment time course. This approach assessed the response to two chemotherapies (cisplatin, paclitaxel) and two metabolic inhibitors (sodium cyanide, 2-deoxy-glucose) across two colorectal PCO lines. The quantitative image analysis pipeline tracked drug response in each PCO before treatment and over multiple time points post-treatment up to 48 hours. Bivariate analysis revealed that morphological, redox ratio, and autofluorescence intensity measurements of drug response provide complementary information. Linear mixed-effect models were used to analyze these longitudinal organoid-level data, which provided improved sensitivity to drug response compared to conventional methods that pool organoid response over time (i.e., do not track individual PCO response). A multivariate analysis of organoid-level redox ratio, autofluorescence intensity, and morphological variables showed distinct sub-populations of PCOs pre-treatment, and heterogeneity in treatment-induced changes across PCOs. This work demonstrates that widefield one-photon redox imaging is an accessible tool for monitoring changes in morphology and metabolism in PCOs, and that single-organoid tracking provides improved sensitivity to PCO drug response.

## 2 Methods

### 2.1 Patient-Derived Cancer Organoid Culture

PCOs were generated from two patient-derived metastatic colorectal cancer lines (P1 and P2), according to a previously published protocol.^35^ Five wells (one well for each of the five treatment groups) were generated in each of the 24-well plates. The base culture medium used was DMEM/F12 (ThermoFisher) supplemented with 10% FBS (Sigma), and 1% Penicillin-Streptomycin (Sigma). Briefly, previously grown PCOs were singularized with 0.25% trypsin, resuspended in base medium, and mixed with Matrigel (Corning # 354234) at a 1:1 ratio. The cell-Matrigel mixture was pipetted onto the glass surface of each well in a 24-well glass bottom plate (black frame, #0 cover glass, Cellvis, P24-0-N). The plate was incubated at 37°C for 2-3 minutes to allow the mixture to solidify and then inverted to ensure PCOs were suspended in the Matrigel. The mixture was left to solidify for at least 20 minutes, then base medium supplemented with Wnt3a-conditioned medium (1:1 ratio) in each well (Fig. 1A, Organoid Culture). Medium was refreshed every two days. Aberrant Wnt/β-catenin signaling is a hallmark of colorectal cancer, and previous studies show that medium conditioned with Wnt3a supports colorectal PCO growth.^36^ Wnt3a-conditioned medium is generated by harvesting medium in which murine L Wnt3a cells (ATCC, CRL-2647) have been cultured.

**Figure 1.**
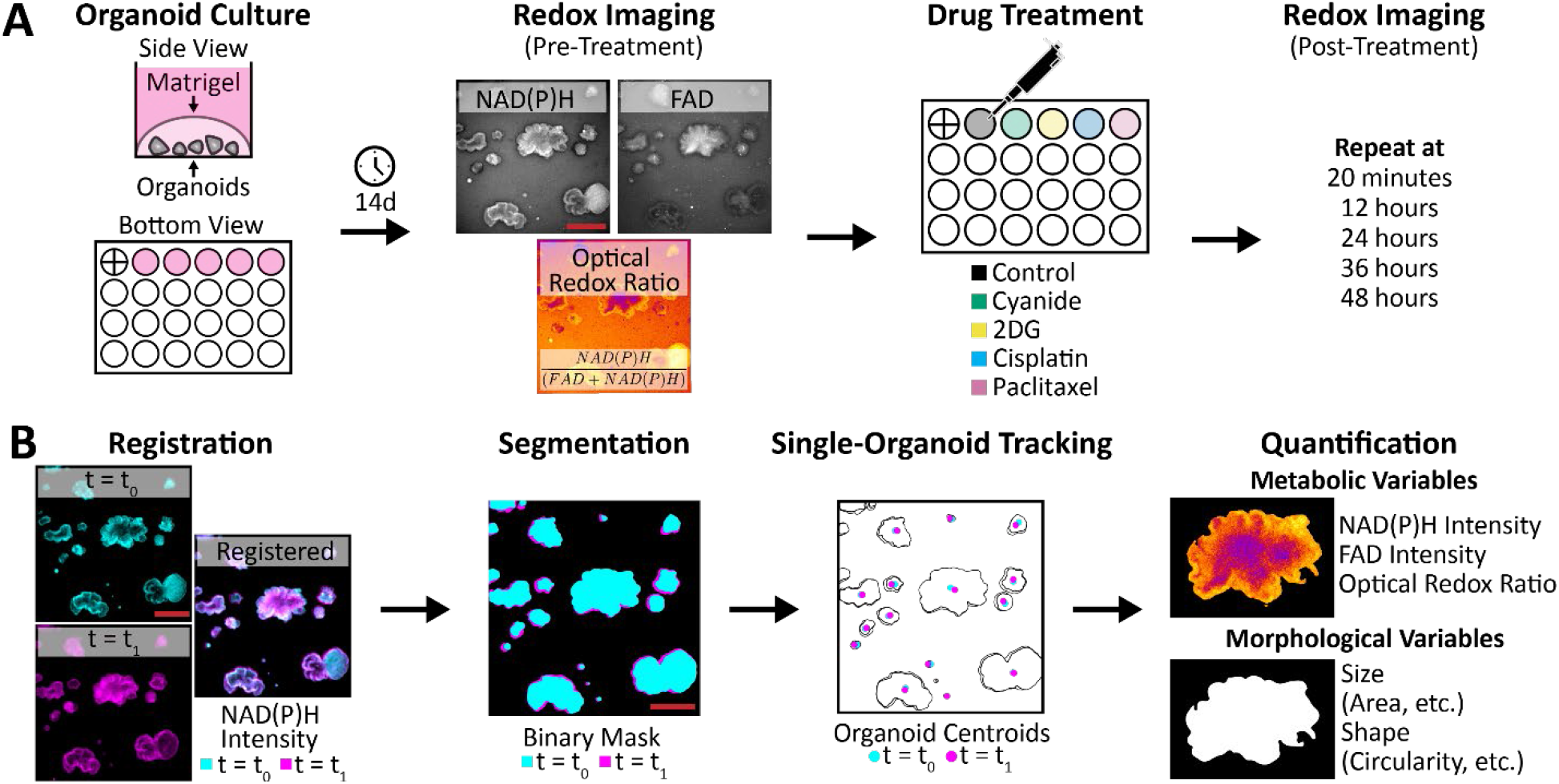
Redox Imaging of Patient-Derived Cancer Organoids. An overview of the protocol for redox imaging and quantitative image analysis. (A) Graphical protocol showing the culturing of patient-derived cancer organoids, pre-treatment redox imaging, drug treatment, and post-treatment redox imaging time course. Redox imaging uses pairs of NAD(P)H and FAD fluorescence images from the same field-of-view to calculate the optical redox ratio image [NAD(P)H/(FAD+NAD(P)H)]. “+” in first well indicates registration mark. (B) Graphical representation of the image analysis pipeline, which includes registration of all frames in each image time series, organoid segmentation, single-organoid tracking, and quantification of metabolic and morphological variables to capture patient-derived cancer organoid drug response at each time point. Scale bar: 500μm.

### 2.2 Treatment Protocol

Five wells for each PCO line were grown for 14 days before treatment. At day 14, media in the wells were refreshed with normal growth medium or medium containing sodium cyanide, 2-deoxy-glucose (2DG), cisplatin, or paclitaxel (Fig. 1A, Drug Treatment). Specifics of each treatment group, including the mechanism of action, concentration used, and number of PCOs per group, are listed in Table 1.

**Table 1.**
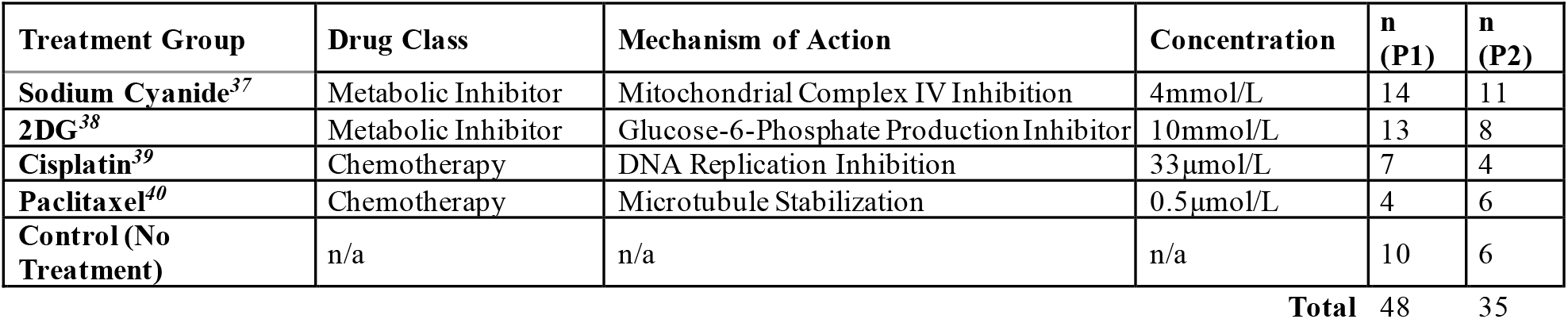
Treatment groups, drug concentrations, and number of PCOs (n) assessed for P1 and P2 lines.

| Treatment Group | Drug Class | Mechanism of Action | Concentration | n (P1) | n (P2) |
| --- | --- | --- | --- | --- | --- |
| **Sodium Cyanide**<sup>37</sup> | Metabolic Inhibitor | Mitochondrial Complex IV Inhibition | 4mmol/L | 14 | 11 |
| **2DG**<sup>38</sup> | Metabolic Inhibitor | Glucose-6-Phosphate Production Inhibitor | 10mmol/L | 13 | 8 |
| **Cisplatin**<sup>39</sup> | Chemotherapy | DNA Replication Inhibition | 33μmol/L | 7 | 4 |
| **Paclitaxel**<sup>40</sup> | Chemotherapy | Microtubule Stabilization | 0.5μmol/L | 4 | 6 |
| **Control (No Treatment)** | n/a | n/a | n/a | 10 | 6 |

**Total** 48 35

### 2.3 Redox Imaging

Redox imaging was performed using an inverted epifluorescence microscope (Nikon Ti-U). System specifications included a 4X air objective (Nikon CFI Plan Fluor, NA 0.13, FOV: 3.33mm x 3.33mm, lateral resolution: 2.16μm at 460nm/2.46μm at 525nm), a white light LED source (SOLA FISH, Lumencor), a scientific grade CMOS camera (Flash4, Hamamatsu, image pixel size: 1.625μm), and an XYZ automated stage (MS-2000, ASI). Microscope control was performed with μManager.^41^ NAD(P)H fluorescence was excited with a DAPI filter cube (Nikon, ex: 361-389nm/em: 435-485nm) and integrated over 3s. FAD fluorescence was excited with a GFP filter cube (Nikon, ex: 450-490nm/em: 500-550nm) and integrated over 5s.

An NAD(P)H fluorescence image and FAD fluorescence image were acquired for each well pre-treatment and at 20 minutes, 12 hours, 24 hours, 36 hours, and 48 hours post-treatment (Fig. 1A, Redox Imaging). The NAD(P)H and FAD images were used to calculate the 2D optical redox ratio for each pixel position (i,j) using Equation 1:

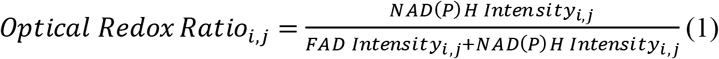

The entire imaging protocol including XYZ stage repositioning and image acquisition was automated with μManager. For each plate, a total of five wells were imaged. Total imaging time per plate was less than 2 minutes, capturing 48 PCOs across five wells for P1 and 35 PCOs across five wells for P2. Note that the first well of the 24-well plate was used to assist in positioning the plate before imaging, which enabled reproducible measurements of the same field-of-view over multiple time-points (Fig. 2A).

**Figure 2.**
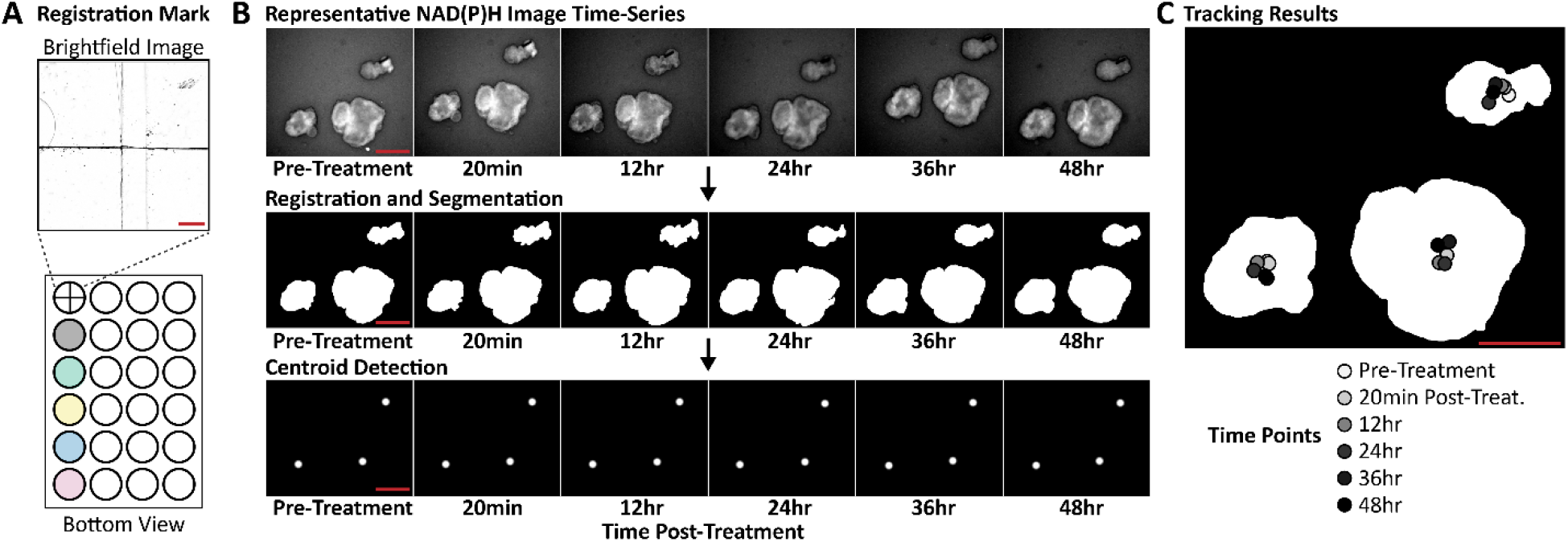
Single-Organoid Tracking. Organoid tracking over multiple time-points was achieved with a registration mark and single-particle tracking. (A) A registration mark was inscribed on the bottom of the first well of the 24-well plate and referenced at each time-point to ensure accurate XYZ positioning. (B) Organoid tracking was achieved by registering the NAD(P)H image time series, segmenting the organoids in each frame, and detecting the centroids of each segmented organoid. These centroids were tracked using an open-source single-particle tracking tool (TrackMate). (C) A segmented organoid mask at 48 hours post-treatment is overlaid with the centroid tracking results at each imaging time-point. Scale bar: 500μm.

### 2.4 Quantitative Image Analysis

A framework for redox imaging and automated image analysis was developed to track each PCO across multiple time points and quantify organoid metabolism and morphology (Fig. 1B). Linkage of each organoid over time was performed via a combined approach of NAD(P)H image time series registration and a 2D centroid matching-based approach with single-particle tracking. This pipeline was implemented using MATLAB (2019b, Mathworks), except when specifically noted.

Registration of each image time series was achieved using both a registration mark and rigid image registration (Fig. 2A,B). A registration mark was etched on the glass bottom of the first well of each plate using a diamond-tip scribe and was referenced at every time point to precisely position the plate before imaging to minimize XYZ drift (Fig. 2A). After acquisition, XY drift between frames was corrected through rigid image registration, which registers two frames by maximizing the cross-correlation of pixel values.^42,43^ The NAD(P)H image time series was used to calculate the XY shift values needed to align the frames in the NAD(P)H image time series, and these shift values were applied to the corresponding FAD and optical redox ratio image time series. These shift values were also used to determine areas that are common across all frames. Image areas that did not overlap across the image time series were excluded.

Segmentation of the PCOs was independently performed on the NAD(P)H image at each time point to capture changes in organoid size and shape (Fig. 2B). However, segmentation based on intensity alone is challenging because autofluorescence images have a low signal-to-background ratio (typically between 2 and 3). A segmentation algorithm based on edge enhancement was developed that reliably segmented all PCOs imaged in this study. The algorithm is as follows:

1. Median filter was applied to remove noise *(medfilt2,* kernel size: 25×25 pixels).
2. Gaussian filter was applied to estimate the background, and this background was subtracted from the image (*imgaussfilt*, kernel size: 450×450 pixels).
3. Local standard deviation filter was applied to enhance edges *(stdfilt,* kernel: 13×13 pixels).
4. Otsu’s method was used to segment the edge-enhanced image into three classes *(imquantize, multithresh,* number of threshold values: 2). The lowest class, corresponding to the non-organoid regions, was excluded.
5. Segmented regions less than 100 pixels were removed.
6. Holes in segmented regions were filled in *(imfill,* option: ‘holes’).
7. Segmented regions were eroded with a disk structure element (*imerode*, *strel*, options: ‘disk’, 9 pixels).
8. Active contours refined the segmented regions *(activecontours,* options: ‘Chan-Vese’, 200 iterations, −0.6 contraction bias).
9. Separation of touching organoids in the mask was achieved via the standard h-Minima-Watershed Transform algorithm.
10. Holes in segmented regions were filled-in *(imfill,* option: ‘holes’).
11. Edges in segmented regions were smoothed with a Gaussian filter (*imgaussfilt*, kernel size: 5×5 pixels).
12. Segmented regions corresponding to PCOs were filtered based on morphology *(regionprops, ismember,* options: regions with area > 1000 pixels and circularity > 0.4).^44^

Specific values for algorithm parameters were chosen empirically based on visual inspection. Validation of the segmentation algorithm was performed by calculating the Sørenson-Dice similarity coefficient for pairs of automatically segmented images and manual segmented images (Figure S1, n = 5, mean ± SEM = 0.93 ± 0.05).

Tracking of each PCO was achieved using an open-source single-particle tracking tool *(TrackMate,* ImageJ/FIJI).^45^ TrackMate achieves single-particle tracking by detecting particles in each frame, linking detected particles from frame-to-frame, and combining all links into the most likely tracks.^46^ Image time series were generated using MATLAB with 2D Gaussian particles (kernel: 10×10 pixels) at the centroid of each segmented PCO (Fig. 2B, Centroid Detection). These centroids were then identified in each frame using the difference-of-Gaussian blob detector (options: 10 pixel diameter, no spot filtering). The linear assignment problem (LAP) algorithm was chosen to perform particle tracking because it is sensitive to events such as particle drift, particle merging, or over-/under-segmentation (options: 200 pixel maximum linking distance, 200 pixel max gap-closing distance, 2 frame max gap-closing distance). All key parameters were initially chosen by visual inspection of the tracking results for the P1 control time series, and then applied to all other image time series (Fig. 2C). Once completed for an image time series, the single-organoid tracking results were used to link the metabolic and morphological variables for each organoid over time.

Quantification of variables that describe organoid morphology and metabolism was performed for each PCO at every time point (n = 83 PCOs total, n = 48 PCOs for P1, n = 35 PCOs for P2). In total, 24 variables (12 metabolic, 12 morphological) were quantified for each segmented organoid: mean, minimum, maximum, and standard deviation of the optical redox ratio, NAD(P)H intensity, and FAD intensity values; organoid area, perimeter, solidity, extent, eccentricity, circularity, minimum Feret’s diameter, maximum Feret’s diameter, minor axis, major axis, convex area, and equivalent diameter *(regionprops).* Detailed definitions of these variables are found in Table S1. PCOs connected to the image border were excluded from statistical analysis.

### 2.5 Statistics

All statistical analyses were performed using MATLAB or R. Open-source MATLAB toolboxes used were Gramm, a data visualization and statistics package, and drtoolbox, a dimensionality reduction package.^47,48^ Open-source R toolboxes included nlme, a mixed effect modeling package, and lsmeans, a least-squares means estimation package.^49,50^

Linear regression was also used to assess the bivariate relationship between organoid area and optical redox ratio, NAD(P)H intensity, and FAD intensity pre-treatment *(stat_glm,* gramm).^48^ The linear pairwise correlations were also calculated for all 24 variables pre-treatment (*corr*).

Organoid-level-normalized time series were calculated for each organoid as X_post-treatment_/X_pre-treatment organoid_, where X is the mean optical redox ratio or organoid area. Linear mixed-effect models were used to analyze these organoid-level-normalized time series for each condition and PCO line *(nlme,* R).^49^ Linear mixed-effect models can account for organoid-level differences in measurements using a random-intercept model, which provides a way to assess treatment-induced effects in the presence of organoid-level heterogeneity.^51,52^ A random-intercept model was specified as *Y*~ *Time* + *Treatment* + *Treatment* * *Time* + (1| *Organoid)* + *Error*, where Y is either the pre-treatment-normalized optical redox ratio or organoid area, Time is a categorical variable encoding time, Treatment is a categorical variable encoding the five treatment groups, Organoid is the organoid-level-normalized value at 20 minutes post-treatment, and Error is the random error in the model. Time is defined as a categorical variable to avoid the assumption of linearity between time and the optical redox ratio or organoid area. Organoid here is encoded as a random effect to account for variability at the organoid-level. Pre-treatment values were excluded from the analysis because all time points were normalized to pre-treatment values (i.e., pretreatment variance is one). A first-order autoregressive (i.e., AR(1)) covariance structure was used to account for the correlations found in these time-course data, where the correlation between time points decreases with increasing separation in time. Pairwise comparisons were performed between all treatment groups at each time point using least-squares means *(lsmeans,* R).^50^ Tukey’s Honest Significant Difference method was used to account for multiple comparisons.

Comparison between single-organoid tracking and pooled analysis was performed. The pooled analysis was the same as the single-organoid tracking analysis, except for the normalization approach and statistical model. To mimic pooled organoid data without organoid tracking, data for each organoid was normalized to the well-level pre-treatment mean as X_post-treatment_/X_pre-treatment well_, where X is the mean optical redox ratio or organoid area. A fixed effects model was specified as *Y~ Time* + *Treatment* + *Treatment* * *Time* + *Error*, where Y is either the well-level-normalized optical redox ratio or organoid area, Time is a categorical variable encoding time, Treatment is a categorical variable encoding the five treatment groups, and Error is the random error in the model. Pre-treatment values were also not included in the analysis. All other parts of the pooled analysis were the same as the organoid-level analysis including the AR(1) covariance structure, least-squares mean for pairwise comparisons, and correction for multiple comparisons.

Principal component analysis (PCA) was used to explore the multivariate relationships between the 24 variables measured from each PCO. PCA was performed using an open-source MATLAB toolbox for dimensionality reduction (*compute_mapping*, *out_of_sample*, drtoolbox). Data from all PCOs pre-treatment, 24 hours post-treatment, and 48 hours post-treatment were standardized within each time point before PCA to account for differences in the scale of each variable *(zscore).* The loadings that projected the data into the principal component (PC) basis were computed using the data from all PCOs pre-treatment. These loadings were then applied to data at 24 hours and 48 hours post-treatment to visualize the time-dependent effects of treatment on the variables from each PCO. Loading vectors were plotted to visualize the contribution of each variable to the PCs. For clarity, only the top 12 variables that loaded on PC1 and PC2 were displayed as they are the variables that contributed the most variability. The projected data were scaled to fit in the loadings interval *(biplot).* The scaled scores of the data projected on the first two PCs were plotted and fitted with ellipses (i.e., bivariate normal distribution) that contained 95% of the points for each group (*geom_point*, *stat_ellipse*, gramm).

## 3 Results

### 3.1 Optical Redox Ratio is Independent of Organoid Morphology

The goal of this study was to segment and track individual PCOs with widefield one-photon redox imaging to extract morphological and metabolic variables of drug response. A quantitative image analysis framework was developed to automatically quantify 12 metabolic variables that summarize NAD(P)H intensity, FAD intensity, and optical redox ratio, along with 12 morphological variables that quantify organoid size and shape (Table S1), for each organoid over time. Relationships between the area of a PCO and its mean NAD(P)H intensity, FAD intensity, and optical redox ratio are shown in Fig. 3A,B,C. Linear models were fit to paired measurements from all PCO pre-treatment (n = 48 PCOs for P1, n = 35 PCOs for P2). There is no correlation between the mean optical redox ratio of each PCO and its area (Fig. 3A, r = 0.04, p = 0.70). However, there is a moderate positive correlation between organoid area and mean NAD(P)H intensity (Fig. 3B, r = 0.48, p < 0.001) and between organoid area and mean FAD intensity (Fig. 3C, r = 0.36, p < 0.001).

**Figure 3.**
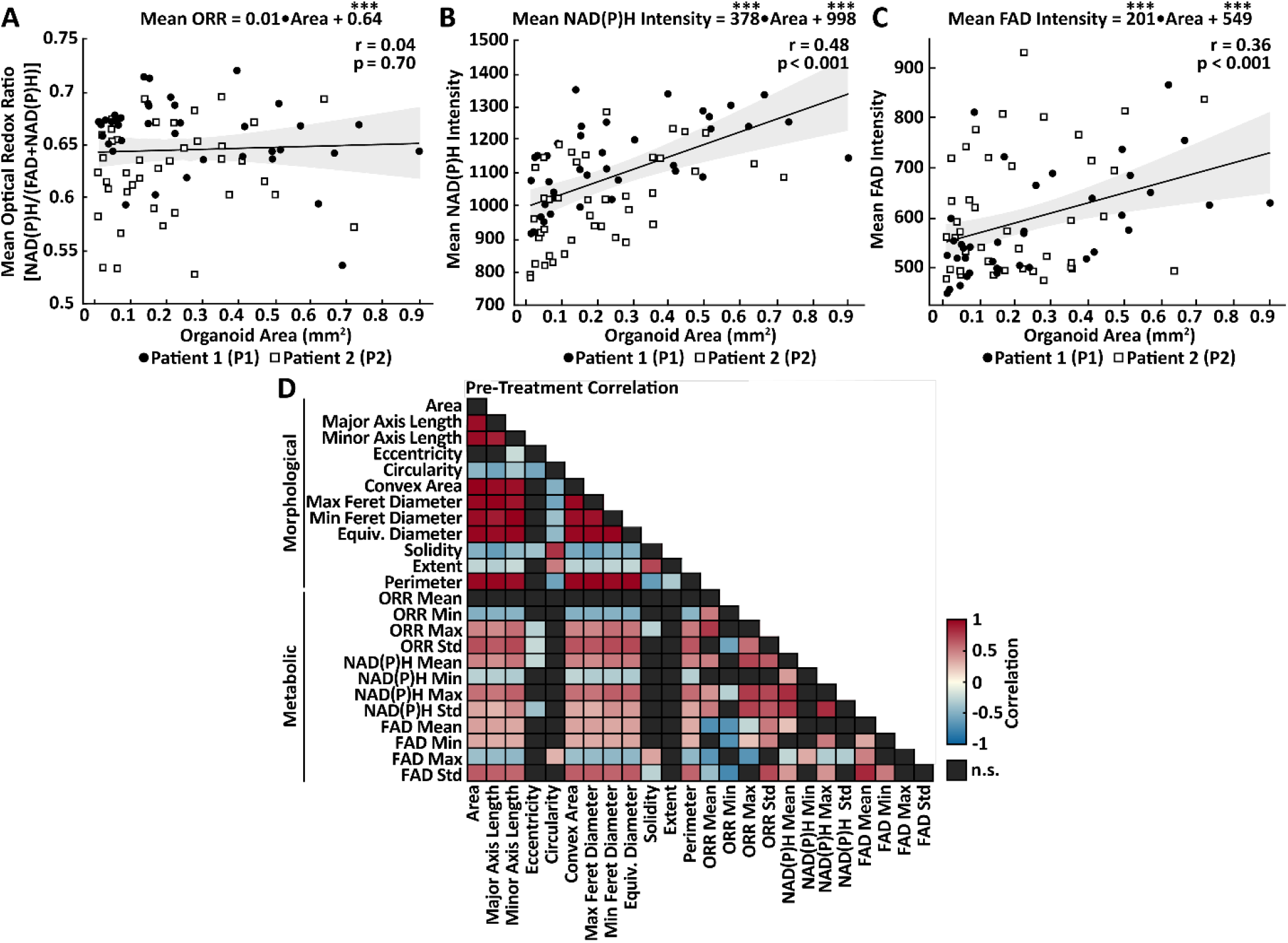
Analysis of Organoid-Level Redox Imaging Variables. Analysis of variables extracted from all patient-derived cancer organoids pre-treatment. (A) Linear regression was used to quantify the relationship between organoid area and its mean optical redox ratio. The fitted model indicates the mean optical redox ratio (ORR) value of each patient-derived cancer organoid is independent of its area (r=0.04, p = 0.70) for all patient-derived cancer organoids from Patient 1 (P1) and Patient (P2) pre-treatment. Statistical significance of the model coefficients is indicated as * * * for p < 0.001. (B) Linear regression was used to quantify the relationship between organoid area and its mean NAD(P)H intensity. The fitted model indicates a moderate positive correlation (r = 0.48, p < 0.001) between organoid area and mean NAD(P)H intensity. (C) Linear regression was used to quantify the relationship between organoid area and its mean FAD intensity. The fitted model indicates a moderate positive correlation (r= 0.36, p < 0.001) between organoid area and mean FAD intensity. (D) Linear pairwise correlation (Pearson’s *r*) matrix of the variables extracted from each patient-derived cancer organoids. Heatmap shows only statistically significant correlations (p < 0.05); n.s., not significant (p > 0.05, black boxes). Table 1 shows the number of patient-derived cancer organoids in each treatment group for each patient.

The linear pairwise correlations of all 24 variables were computed to understand the relationships between the metabolic and morphological variables (Fig. 3D). As expected, morphological variables were significantly correlated (p < 0.05) with most other morphological variables apart from eccentricity, which was only significantly correlated with the minor axis length, solidity, and circularity. Likewise, metabolic variables were significantly correlated (p < 0.05) with many other metabolic variables except for the minimum and maximum values, which were uncorrelated with many of the other metabolic variables. Statistically significant correlations between morphological and metabolic variables were also observed, especially for standard deviations in FAD intensity and optical redox ratio that positively correlate with greater organoid area/diameter/length. Consistent with the linear model (Fig. 3A), the mean optical redox ratio had no significant correlations with any morphological parameters. Previous studies have shown that the optical redox ratio of cancer organoids predicts treatment response^9–12^, so this data indicates that the mean optical redox ratio may provide information about PCO drug response that is independent of morphology.

### 3.2 Drug Response Lowers Optical Redox Ratio

Changes in the optical redox ratio were calculated in two PCO lines over a 48 hour treatment time course. Previous studies have found that a treatment-induced decrease in the optical redox ratio of cancer organoids predicts *in vivo* treatment response.^23,29^ Treatment-induced changes in mean NAD(P)H intensity and mean FAD intensity can be found in Figure S2A,B,C,D. Both PCO lines were treated with cisplatin and paclitaxel, two chemotherapies, and sodium cyanide and 2DG, two metabolic inhibitors. There was one well for each treatment and an additional untreated well served as a negative control (five wells total). Three representative optical redox ratio image time series are shown in Figure 4A, which show the change in optical redox ratio with cyanide treatment, large change in optical redox ratio with paclitaxel treatment, and the stability of the optical redox ratio in untreated organoids. Cyanide and paclitaxel treatment were representative of a minimal and large response, respectively. Arrows in Figure 4A indicate specific PCOs that are representative of changes observed for that condition. Single-organoid tracking was used to normalize each PCO to its pre-treatment optical redox ratio, so that relative changes in optical redox ratio could be compared across PCOs. Least-squares means was used to estimate the mean of each group, and to calculate the pairwise differences at each time point between treatment groups (Fig. 4B,C).

**Figure 4.**
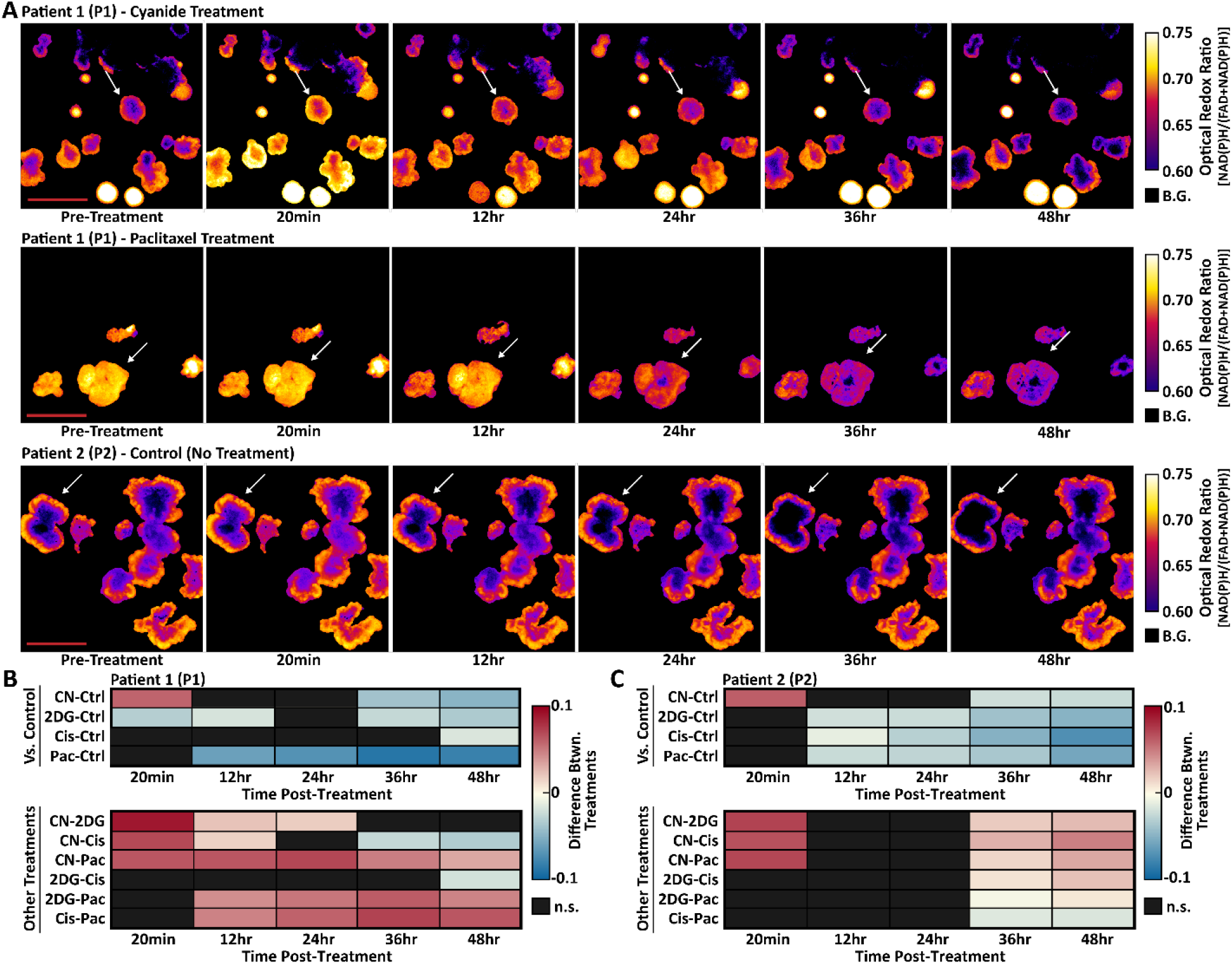
Treatment-Induced Changes in Patient-Derived Cancer Organoid Optical Redox Ratio. Quantitative im a ge analy sis tracked changes in the optical redox ratio of each patient-derived cancer organoid over time. (A) Representative optical redox ratio image time series for Patient 1 (P1) and Patient 2 (P2). Arrows indicate specific organoids representative of changes observed for that condition. B.G, background. (B) Heatmaps show the pairwise percent differences between pre-treatment-normalized optical redox ratio between all treatment groups for P1. Top: Pairwise percent differences between each treatment and control for P1. Bottom: Pairwise percent differences between drug treatments (excluding control) for P1. (C) Heatmaps show the pairwise percent differences between pre-treatment-normalized optical redox ratio between all treatment groups for P2. Top: Pairwise percent differences between each treatment and control for P2. Bottom: Pairwise percent differences between drug treatments for P2. All pairwise differences were calculated from the linear mixed-effect models via least-squares means. n.s., not significant (p > 0.05, black boxes). Table 1 shows the number of patient-derived cancer organoids in each treatment group for each patient. Scale bar: 500μm.

For P1 PCOs, the optical redox ratio decreased compared to control at 36 hours for the cyanide (p < 0.01), 2DG (p < 0.01) groups (Fig. 4B). Cisplatin treatment in P1 resulted in a decreased optical redox ratio compared to control only at 48 hours (Fig. 4B, p < 0.05). Paclitaxel treatment in P1 resulted in a decreased optical redox ratio compared to control starting at 12 hours (Fig 4B, p < 0.05). For P2 PCOs, the optical redox ratio decreased compared to control at 12 hours for the 2DG (p < 0.01), cisplatin (p < 0.05), and paclitaxel (p < 0.05) groups (Fig. 4C). Cyanide treatment in P2 resulted in a decreased optical redox ratio compared to control starting at 36 hours (Fig. 4C, p < 0.01). Additionally, cyanide in both P1 and P2 caused a sharp rise in the optical redox ratio (p < 0.001) at 20 minutes, characteristic of electron transport chain inhibition (Fig. 4B,C). Heatmaps (Fig. 4B,C) also show the pairwise differences between treatment groups.

### 3.3 Drug Response Slows Organoid Growth

Changes in organoid size or growth rate can occur due to effective drug response, so the relative change in organoid area over time was also quantified. All other morphological variables remained constant or had a limited change over 48 hours compared to pre-treatment values (a subset can be found in Fig. S2E-J). Three representative image time series in Figure 5A showing segmented PCOs with the color-coded value mapped to the pre-treatment normalized organoid area. These image time series illustrate increased organoid area over time in control conditions, and relatively modest increases in organoid area over time in cyanide and paclitaxel treatment conditions. Arrows in Figure 5A indicate specific PCOs representative of changes observed for that condition. Heterogeneity in organoid area was observed pre-treatment, so single-organoid tracking was used to normalize each organoid to its pre-treatment area. This enabled comparisons of relative changes in area across organoids. Least-squares means was used to estimate the mean of each group, and to calculate the pairwise differences at each time point between control and treatment groups (Fig. 5B,C).

**Figure 5.**
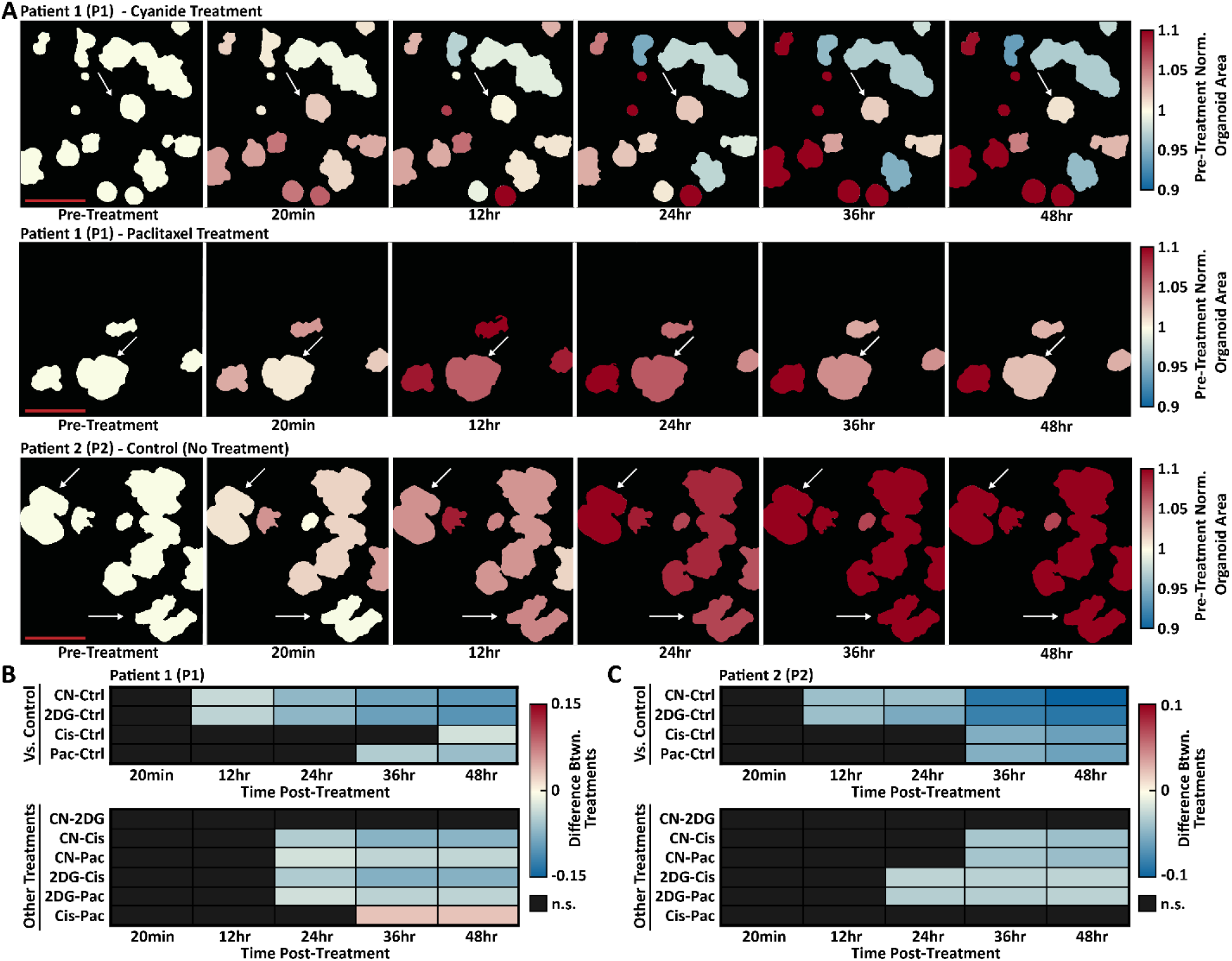
Treatment-induced Changes in Patient-Derived Cancer Organoid Areas. Quantitative im age analy sis tracked changes in the area of each patient-derived cancer organoid over time. (A) Representative segmented image time series for Patient 1 (P1) and Patient 2 (P2). The color-coded value of each segmented organoid corresponds to value at each time-point divided by its pre-treatment value. Arrows indicate specific organoids that are representative of changes observed for that condition. (B) Heatmaps show the pairwise percent differences between pre-treatment-normalized area between all treatment groups for P1. Top: Pairwise percent differences between each treatment and control for P1. Bottom: Pairwise percent differences between drug treatments (excluding control) for P1. (C) Heatmaps show the pairwise percent differences between pre-treatment-normalized area between all treatment groups for P2. Top: Pairwise percent differences between each treatment and control for P2. Bottom: Pairwise percent differences between drug treatments for P2. All pairwise differences were calculated from the linear mixed-effect models via least-squares means. n.s., not significant (p > 0.05, black boxes). Table 1 shows the number of patient-derived cancer organoid in each treatment group for each patient. Scale bar: 500μm.

Two separate patterns of response emerged for metabolic inhibitors and chemotherapies. For P1 groups treated with metabolic inhibitors, lower organoid areas compared to control were observed starting at 12 hours for the cyanide (p < 0.01) and 2DG (p < 0.01) (Fig. 5B). However, paclitaxel-treated P1 PCOs continued to grow until 24 hours and were only significantly different from control beginning at 36 hours (p < 0.05) (Fig. 5B). For P1 PCOs, cisplatin treatment resulted in lower organoid areas compared to control only at 48 hours (p < 0.05) (Fig. 5B). For P2 PCOs treated with metabolic inhibitors, lower organoid areas compared to control were observed starting at 12 hours for the cyanide (p < 0.01) and 2DG (p < 0.01) groups (Fig. 5C). For P2 PCOs treated with chemotherapies, continued growth was observed until 24 hours, and organoid area was only lower than control beginning at 36 hours for paclitaxel (p < 0.01) and cisplatin (p < 0.01) groups (Fig. 5C). Heatmaps (Fig. 5B,C) also show the pairwise differences between treatment groups.

### 3.4 Single-Organoid Tracking Provides Improved Sensitivity

PCOs are heterogeneous and exhibited a range of optical redox ratio and organoid area values. Single-organoid tracking can monitor dynamic changes in organoid-level variables over time by normalizing each PCO to its own pre-treatment value. This enabled the observation of the optical redox ratio and organoid area on an organoid-level in the P1 and P2 control groups. In both P1 and P2 control groups, the optical redox ratio was stable over 48 hours (Fig. 6A-C, p > 0.05 pretreatment vs. each time-point post-treatment using organoid-level normalization). In both P1 and P2 control groups, PCOs continue to grow over 48 hours, with a statistically significant increase in organoid area of 10% and 11%, respectively, at 48 hours compared to pre-treatment (Fig. 6E,G, p < 0.05 pre-treatment vs. each time-point post-treatment using organoid-level normalization).

**Figure 6.**
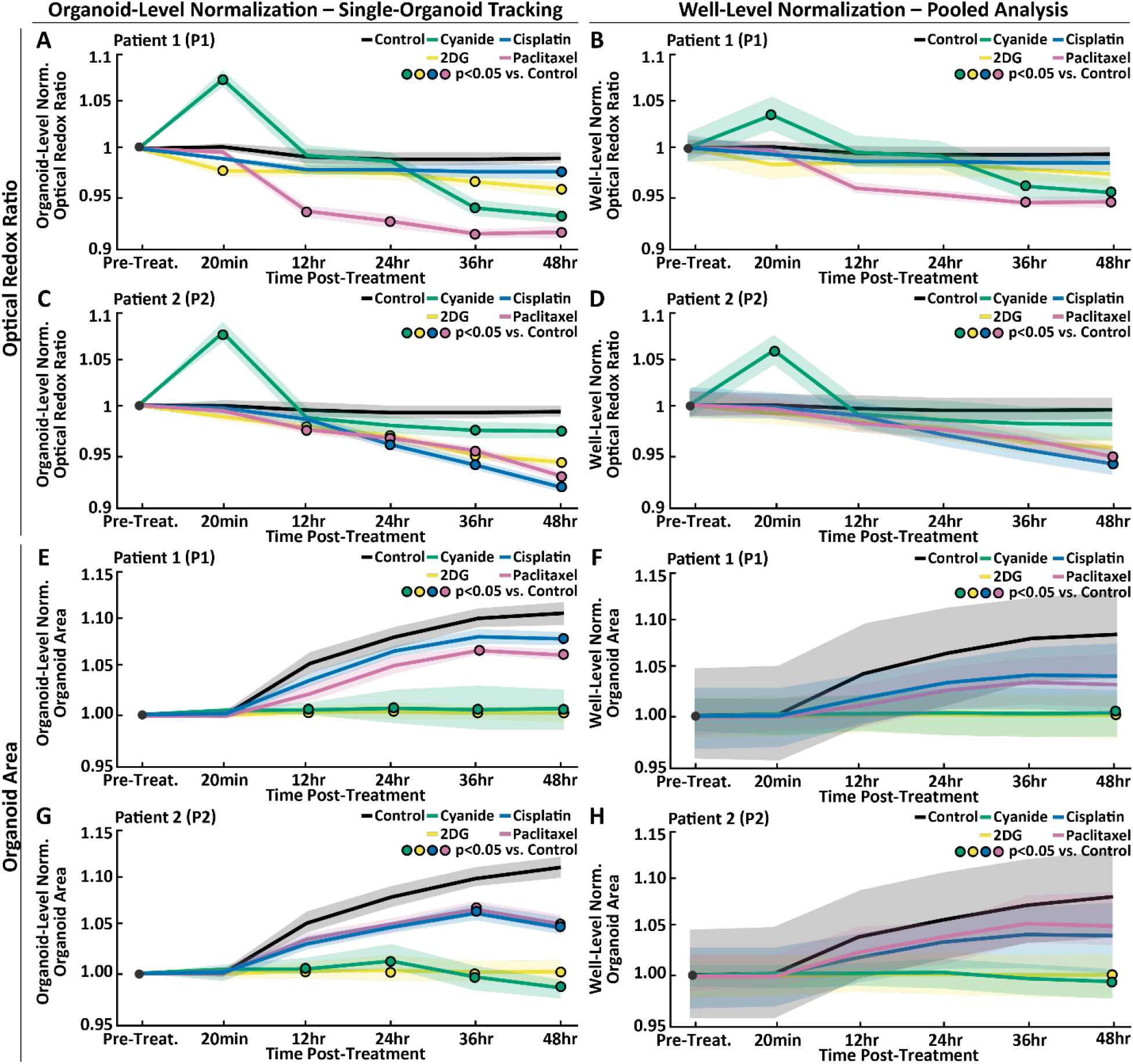
Comparison of Single-Organoid Tracking and Pooled Analysis. Comparison of single-organoid tracking and pooled analysis using time series data from Patient1 (P1) and Patient 2 (P2). Normalization refers to dividing the value of each organoid at each time-point by either its own pre-treatment value (organoid-level normalization) or the well-level mean (well-level normalization). (A,B) Pre-treatment normalized optical redox ratio using organoid-level normalization (left) and well-level normalization (right) for each treatment group for P1. (C,D) Pre-treatment normalized optical redox ratio using organoid-level normalization (left) and well-level normalization (right) for each treatment group for P2. (E,F) Pre-treatment normalized organoid area using organoid-level normalization (left) and well-level normalization (right) for each treatment group for P1. (G,H) Pre-treatment normalized organoid area using organoid-level normalization (left) and well-level normalization (right) for each treatment group for P2. Data in A,C,E,G were analyzed using a linear mixed-effect model that uses single-organoid tracking to account for organoid-level variability. Data in B,D,F,H were analyzed using a linear mixed-effect model that does not account for organoid-level variability due to the lack of single-organoid tracking. Data plotted is mean (lines) ± standard error of the mean (shaded area). Significant differences between a treatment and control (p < 0.05) is indicated with a circle color-coded for the treatment group (see legend). Table 1 shows the number of patient-derived cancer organoids in each group and for each patient.

To demonstrate the utility of single-organoid tracking, a pooled analysis was performed. In addition to the previous analysis using organoid-level normalization, data was analyzed using well-level normalization where each PCO was normalized to the well-level mean. Figure 6 shows the optical redox ratio and organoid area over time for P1 and P2 analyzed using single-organoid tracking (organoid-level normalization) or pooled analysis (well-level normalization). Single-organoid tracking (Fig. 6A,C,E,G) provided data with lower variability compared to pooled analysis (Fig. 6B,D,F,H), as the well-level mean does not fully account for organoid-level heterogeneity. In addition, pooled analysis does not provide organoid-level time series for use in a statistical model. Single-organoid tracking found more differences between control and treatment groups when compared to pooled analysis, which demonstrates the improved sensitivity to treatment response afforded by single-organoid tracking.

### 3.5 Multivariate Analysis Reveals Patient-Derived Cancer Organoid Sub-Populations

Two sub-populations of organoids with distinct morphology (solid and hollow) are present in cultures from P1, while all P2 organoids are morphologically similar (Fig. 7A). The first P1 subpopulation, P1 Solid, were generally dense (highly scattering), irregular in morphology, and had core regions with lower optical redox ratio values than the periphery. The second P1 subpopulation, P1 Hollow, were less dense (lower scattering), had a spherical morphology, and had core regions with higher optical redox ratio values than the outer regions. In addition, the optical redox ratio values of P1 Hollow PCOs were generally higher than P1 Solid PCOs, driven by higher NAD(P)H intensity. P2 appeared to have one type of organoid morphology that was qualitatively similar to P1 Solid: dense, irregular in morphology, and with a core region with a low optical redox ratio compared to the periphery. The differences in the internal structure of P1 Solid and P1 Hollow PCOs were confirmed by confocal microscopy of P1 Solid and P1 Hollow PCOs stained for epithelial cell adhesion molecule (EpCAM, Fig. S3). By qualitative examination, 39 P1 Solid, 9 P1 Hollow, and 35 P2 PCOs were identified.

**Figure 7.**
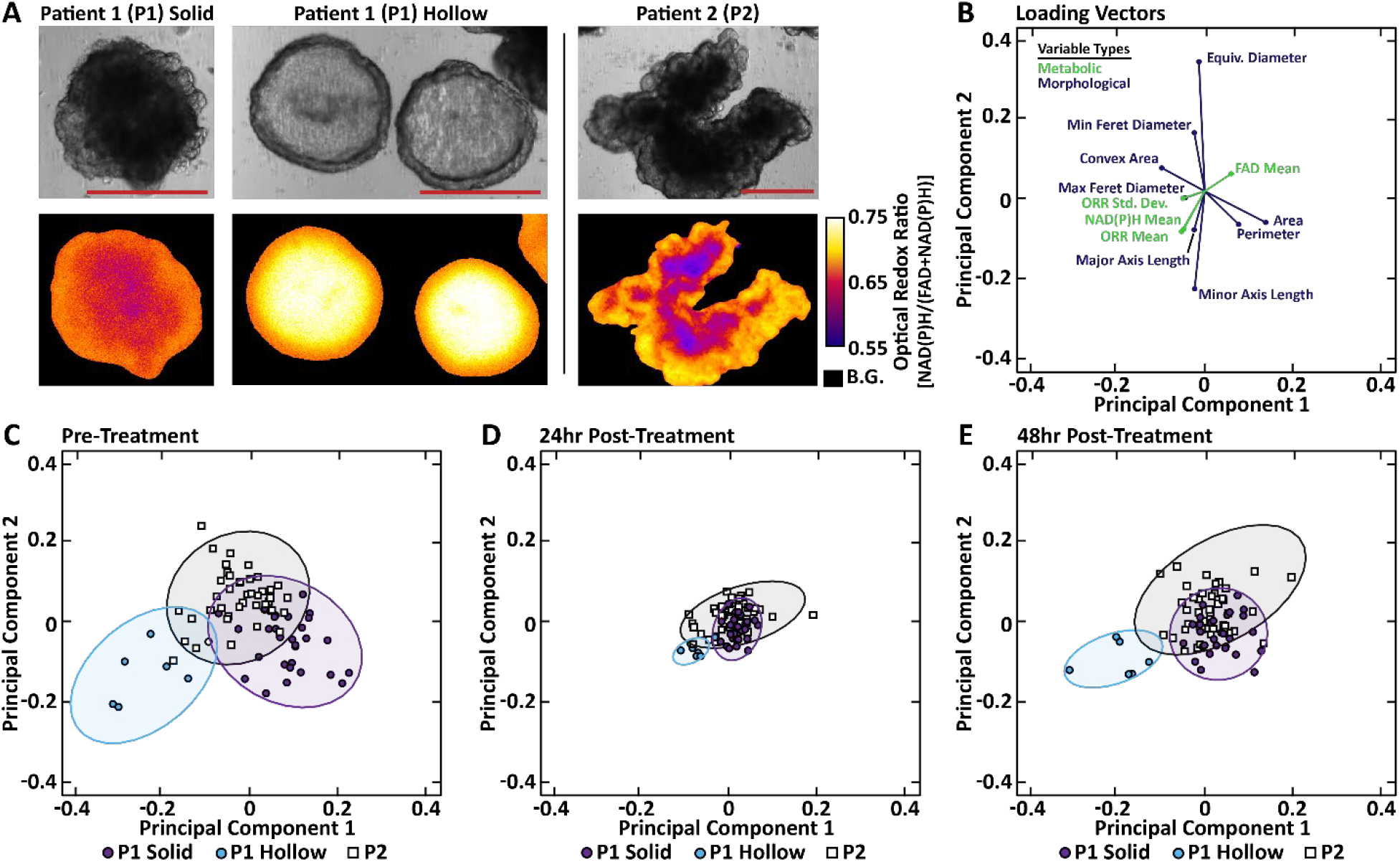
Phenotypic Identification of Patient-Derived Cancer Organoid Sub-Populations. Qualitative and quantitative (principal component analysis or PCA) evaluation of patient-derived cancer organoids with distinct phenotypes from Patient 1 (P1) and Patient 2 (P2). (A) Paired brightfield (top) and optical redox ratio (bottom) images of representative patient-derived cancer organoid from each of the three phenotypic sub-populations: P1 Solid, P1 Hollow and P2. Scale bar: 250μm. B.G., background. (B) Loading vectors for the 12 variables with the highest loading on PC1 and PC2, which were defined with pre-treatment data only. Green are metabolic variables and blue are morphological variables. (C) Data from all organoids projected onto PC1 and PC2. Colors correspond to the three qualitatively identified phenotypic organoid sub-populations: P1 Solid, P1 Hollow, and P2. Ellipse containing 95% of the points for each group were also plotted. (D) The pre-treatment PCs applied to data from the patient-derived cancer organoids at 24 hours post-treatment. (E) The pre-treatment PCs applied to data from patient-derived cancer organoids at 48 hours post-treatment.

Principal component analysis (PCA) was applied to the 24 morphological and metabolic variables (Supplemental Table 1) for all pre-treatment PCOs to determine if these variables could cluster PCOs by P1 Solid, P1 Hollow, and P2. A plot of the loading vectors (Fig. 7B) show the contribution of each of the pre-treatment variables on the first two PCs. The length of the vector corresponds to the magnitude of the contribution of a specific variable to the first two PCs. The angle between vectors, as well as the angle between each vector and the PC axes, indicate the level of correlation. The color indicates which are metabolic or morphological variables. Only the 12 variables that have the largest contributions to the first two PCs were displayed for clarity. Figure 7C shows the organoid-level data projected onto the PC1-PC2 basis, with clear clustering of subpopulations based on P1 Solid, P1 Hollow, and P2. Notably, P1 Hollow was correlated with metabolic variables, clustering in the lower left quadrant apart from the P1 and P2 clusters (Fig. 7B,C). The same pre-treatment PC1-PC2 basis was used to plot PCOs at 24 and 48 hours posttreatment. At 24 hours post-treatment, smaller variability is observed within and between clusters compared to pre-treatment variability (Fig. 7D), and cluster variability increased at 48 hours (Fig. 7E).

## 4. Discussion

Functional drug screens using PCOs, which reflect the heterogeneity and drug sensitivity of *in vivo* tumors, could be used to predict the most effective treatment for a patient and identify novel drug candidates during drug development.^5,6,8,10,12^ However, current methods for evaluating drug response in PCOs are destructive, ignore organoid-level heterogeneity, provide limited metrics of response, or lack throughput. Cell viability assays (e.g., tetrazolium, luciferase) measure metabolic function as a surrogate for the number of viable cells, but use reagents with long-term toxicity, consume the sample, require genetic manipulation, or do not capture heterogeneity.^53–56^ Histology provides cell- and organoid-level structural and molecular information, but requires slow, destructive sample processing.^57^ Standard molecular biology techniques (e.g., RNA-seq, qPCR, Western blots) can capture treatment-induced changes in protein and RNA levels, but are destructive and require the pooling of many PCOs to yield enough cell material.^57^ Organoid diameter measurements from brightfield images can assess the same PCOs over time, but only provide a single measure of response (i.e., change in organoid area), and these measurements are often performed manually, introducing user bias.^11,35^

Here, we provide the first demonstration of widefield one-photon redox imaging for functional assessment of drug response in PCOs. Redox imaging improves on existing tools for PCO screening as it is non-destructive, rapid, and label-free, requiring no additional dyes or reagents. Performing redox imaging in a widefield one-photon epifluorescence configuration is well suited for high-throughput applications because of simple, widely available instrumentation, large field-of-view, micron-scale resolution, and fast acquisition. However, widefield one-photon epifluorescence microscopy lacks optical sectioning, limiting its ability to resolve cellular level detail within scattering samples like organoids. The lack of optical sectioning can also be beneficial for high-throughput PCO screening as it captures an integrated response from the entire organoid and is less sensitive to shifts in axial position. While widefield one-photon redox imaging does not provides a readout at the cell-level, the organoid-level readout may better represent the response of a patient’s tumor as multiple organoids captures more cells for analysis. This study shows that widefield one-photon redox imaging provides automated assessment of multiple quantitative variables at the organoid-level that capture drug response and separate PCO sub-populations based on phenotype. A key result of this study is a framework to acquire robust redox imaging data that links values within the same PCO across multiple time points using a standard epifluorescence microscope. These methods can be adopted by non-experts to achieve reproducible data of drug response in PCOs using accessible instrumentation, while accounting for pre-treatment variability between PCOs to yield meaningful drug response data.

This was the first study to perform widefield one-photon redox imaging of PCOs, so the relationships between the organoid-level morphological and metabolic variables were unknown. The relationship between organoid size and optical redox ratio was of interest as PCOs cultured over long periods of time form a characteristic core of dead cells with a lower optical redox ratio. The lower optical redox ratio is likely due to increased flavin production (riboflavin, flavin mononucleotide, and FAD) during the cellular response to cytotoxic stress, suggesting the optical redox ratio [NAD(P)H/(FAD + NAD(P)H)] is sensitive to cell death.^58^,^59^ Interestingly, we found that the mean optical redox ratio was independent of organoid area for the PCOs imaged in this study (area range: 0.02-0.9mm^2^). Larger PCOs exhibited higher mean FAD intensity values due to larger dead cores, but this was compensated by increased NAD(P)H levels at the periphery, which we hypothesize is due to the presence of more viable cell material. Further analysis of all variables revealed that the mean optical redox ratio is uncorrelated with other measures of organoid morphology, indicating that the optical redox ratio provides complementary information to morphology.

PCOs grow stochastically due to the culture technique (single-cell suspension in a 3D matrix) and cellular heterogeneity, resulting in organoids that are unique in size, morphology, and metabolism pre-treatment and post-treatment.^60,61^ Previous studies of drug response in PCOs have only assessed response at single time point or used different organoids for each time point, both of which only capture a snapshot of the PCO population. Automated single-organoid tracking enables measurement of organoid-level drug response over multiple time points. Tracking organoids over time also overcomes issues with variability in the numbers and sizes of PCOs in each well. Statistical techniques such as mixed-effect effect models can use these time-course data to account for organoid-level heterogeneity and provide improved sensitivity to drug response.^51,62^ Our results show that widefield one-photon redox imaging can determine drug response in PCOs, and that single-organoid tracking improves sensitivity to treatment-induced changes in redox imaging variables (Fig. 6).

PCOs generated from P1 exhibited two distinct phenotypes, P1 Solid and P1 Hollow, which were identified by visual inspection and confirmed by multivariate analysis (PCA). The fluid-filled hollow morphology of P1 Hollow PCOs are consistent with cystic morphologies seen in other papers using colorectal PCOs.^63^ As these phenotypes are stable over multiple passages, these two sub-populations likely reflect the *in vivo* heterogeneity of the patient’s tumor and are not due to mutations gained *in vitro*. The loading vectors (Fig. 7B) appear to confirm that the metabolic and morphological variables are orthogonal, which demonstrates the information gained by redox imaging and is consistent with the limited correlations between organoid metabolism and morphology (Fig. 3D). As the P1 Hollow sub-population comprised only 19% of all P1 PCOs and were not evenly distributed among the five treatment groups, stratification of the response of each sub-population was not attempted.

This study demonstrates a framework for performing widefield one-photon redox imaging of PCOs and single-organoid tracking that accounts for pre-treatment heterogeneity in organoid morphology and metabolism for robust drug response measurements. Widefield one-photon redox imaging is an easily scalable and accessible technology that provides both metabolic and morphological information for characterizing PCO populations. We believe that these validated approaches will enable broad dissemination of widefield one-photon redox imaging for screening of PCOs, with applications in drug development and clinical treatment planning.

## Acknowledgements

We thank Tongcheng Qian for providing the confocal microscopy images. We also thank Tiffany Heaster, Rupsa Datta, Peter Favreau, and Maryse Lapierre-Landry for their valuable feedback on the manuscript. This work was supported by the NSF (CBET-1642287), Stand Up to Cancer (SU2C-AACR-IG-08-16), the NIH (R01 CA185747, R01 CA185747, R37 CA226526) and the University of Wisconsin Carbone Cancer Center (Support Grant P30 CA014520).

## Disclosures

Authors have no relevant financial interests in the manuscript and no other potential conflicts of interest to disclose.

## Notes

### Competing Interest Statement

The authors have declared no competing interest.

## References

1. E. Fountzilas, and A. M. Tsimberidou, “Overview of precision oncology trials: challenges and opportunities,” Expert review of clinical pharmacology 11(8), 797–804 (2018).

2. V. Prasad, “Perspective: The precision-oncology illusion,” Nature 537(7619), S63–S63 (2016).

3. I. F. Tannock, and J. A. Hickman, “Limits to Personalized Cancer Medicine,” New England Journal of Medicine 375(13), 1289–1294 (2016).

4. A. Letai, “Functional precision cancer medicine—moving beyond pure genomics,” Nature Medicine 23(9), 1028–1035 (2017).

5. N. Sachs et al., “A Living Biobank of Breast Cancer Organoids Captures Disease Heterogeneity,” Cell 172(1-2), 373–386 e310 (2018).

6. G. Vlachogiannis et al., “Patient-derived organoids model treatment response of metastatic gastrointestinal cancers,” Science 359(6378), 920–926 (2018).

7. Y. Saito et al., “Establishment of Patient-Derived Organoids and Drug Screening for Biliary Tract Carcinoma,” Cell Rep 27(4), 1265–1276.e1264 (2019).

8. H. Tiriac et al., “Organoid Profiling Identifies Common Responders to Chemotherapy in Pancreatic Cancer,” Cancer Discovery 8(9), 1112 (2018).

9. D. Schumacher et al., “Heterogeneous pathway activation and drug response modelled in colorectal-tumor-derived 3D cultures,” PLOS Genetics 15(3), e1008076 (2019).

10. N. Sasaki, and H. Clevers, “Studying cellular heterogeneity and drug sensitivity in colorectal cancer using organoid technology,” Current Opinion in Genetics & Development 52(117–122 (2018).

11. C. A. Pasch et al., “Patient-derived cancer organoid cultures to predict sensitivity to chemotherapy and radiation,” Clinical Cancer Research clincanres.3590.2018 (2019).

12. O. Kopper et al., “An organoid platform for ovarian cancer captures intra- and interpatient heterogeneity,” Nature Medicine 25(5), 838–849 (2019).

13. J. T. Sharick et al., “Cellular Metabolic Heterogeneity In Vivo Is Recapitulated in Tumor Organoids,” Neoplasia (New York, N.Y.) 21(6), 615–626 (2019).

14. A. Luengo, D. Y. Gui, and M. G. Vander Heiden, “Targeting Metabolism for Cancer Therapy,” Cell Chem Biol 24(9), 1161–1180 (2017).

15. D. A. Gil et al., “Redox imaging and optical coherence tomography of the respiratory ciliated epithelium,” J Biomed Opt 24(1), 1–4 (2019).

16. A. J. Walsh, and M. C. Skala, “Optical metabolic imaging quantifies heterogeneous cell populations,” Biomedical optics express 6(2), 559–573 (2015).

17. A. Walsh et al., “Optical imaging of metabolism in HER2 overexpressing breast cancer cells,” Biomedical optics express 3(1), 75–85 (2012).

18. A. Varone et al., “Endogenous two-photon fluorescence imaging elucidates metabolic changes related to enhanced glycolysis and glutamine consumption in precancerous epithelial tissues,” Cancer Res 74(11), 3067–3075 (2014).

19. M. C. Skala et al., “In vivo multiphoton microscopy of NADH and FAD redox states, fluorescence lifetimes, and cellular morphology in precancerous epithelia,” Proc Natl AcadSci USA 104(49), 19494–19499 (2007).

20. M. C. Skala et al., “Longitudinal optical imaging of tumor metabolism and hemodynamics,” J Biomed Opt 15(1), 011112 (2010).

21. M. Skala, and N. Ramanujam, “Multiphoton redox ratio imaging for metabolic monitoring in vivo,” Methods Mol Biol 594(155–162 (2010).

22. A. T. Shah et al., “In Vivo Autofluorescence Imaging of Tumor Heterogeneity in Response to Treatment,” Neoplasia 17(12), 862–870 (2015).

23. A. T. Shah et al., “Optical Metabolic Imaging of Treatment Response in Human Head and Neck Squamous Cell Carcinoma,” PLOS ONE 9(3), e90746 (2014).

24. A. A. Heikal, “Intracellular coenzymes as natural biomarkers for metabolic activities and mitochondrial anomalies,” Biomark Med4(2), 241–263 (2010).

25. O. I. Kolenc, and K. P. Quinn, “Evaluating Cell Metabolism Through Autofluorescence Imaging of NAD(P)H and FAD,” Antioxidants & Redox Signaling30(6), 875–889 (2017).

26. I. Georgakoudi, and K. P. Quinn, “Optical imaging using endogenous contrast to assess metabolic state,” Annu Rev Biomed Eng 14(351–367 (2012).

27. K. P. Quinn et al., “Quantitative metabolic imaging using endogenous fluorescence to detect stem cell differentiation,” SciRep 3(3432 (2013).

28. A. T. Shah, T. M. Heaster, and M. C. Skala, “Metabolic Imaging of Head and Neck Cancer Organoids,” PLoS One 12(1), e0170415 (2017).

29. T. M. Cannon et al., “High-throughput measurements of the optical redox ratio using a commercial microplate reader,” J Biomed Opt 20(1), 010503 (2015).

30. A. J. Walsh et al., “Optical metabolic imaging identifies glycolytic levels, subtypes, and early-treatment response in breast cancer,” Cancer Res 73(20), 6164–6174 (2013).

31. J. T. Sharick et al., “Metabolic Heterogeneity in Patient Tumor-Derived Organoids by Primary Site and Drug Treatment,” Frontiers in Oncology 10(553), (2020).

32. A. J. Walsh et al., “Quantitative optical imaging of primary tumor organoid metabolism predicts drug response in breast cancer,” Cancer Res 74(18), 5184–5194 (2014).

33. P. F. Favreau et al., “Label-free redox imaging of patient-derived organoids using selective plane illumination microscopy,” Biomedical optics express 11(5), 2591–2606 (2020).

34. N. Phan et al., “A simple high-throughput approach identifies actionable drug sensitivities in patient-derived tumor organoids,” Communications Biology 2(1), 78 (2019).

35. S. L. Fricke et al., “MTORC1/2 Inhibition as a Therapeutic Strategy for PIK3CA Mutant Cancers,” Mol Cancer Ther 18(2), 346–355 (2019).

36. T. Zhan, N. Rindtorff, and M. Boutros, “Wnt signaling in cancer,” Oncogene 36(11), 1461–1473 (2017).

37. J. L. Way, “Cyanide intoxication and its mechanism of antagonism,” Annu Rev Pharmacol Toxicol 24(451–481 (1984).

38. D. Zhang et al., “2-Deoxy-D-glucose targeting of glucose metabolism in cancer cells as a potential therapy,” Cancer Letters 355(2), 176–183 (2014).

39. S. Dasari, and P. B. Tchounwou, “Cisplatin in cancer therapy: molecular mechanisms of action,” Eur J Pharmacol 740(364–378 (2014).

40. B. A. Weaver, “How Taxol/paclitaxel kills cancer cells,” Mol Biol Cell 25(18), 2677–2681 (2014).

41. A. D. Edelstein et al., “Advanced methods of microscope control using μManager software,” J Biol Methods 1(2), e10 (2014).

42. M. Guizar-Sicairos, S. T. Thurman, and J. R. Fienup, “Efficient subpixel image registration algorithms,” Optics letters 33(2), 156–158 (2008).

43. M. Guizar, “Efficient subpixel image registration by cross-correlation,” MATLAB Central File Exchange. (2020).

44. P. Soille, Morphological image analysis: principles and applications, 2nd ed., Springer, Berlin; New York (2003).

45. J. Y. Tinevez et al., “TrackMate: An open and extensible platform for single-particle tracking,” Methods 115(80–90 (2017).

46. K. Jaqaman et al., “Robust single-particle tracking in live-cell time-lapse sequences,” Nature Methods 5(8), 695–702 (2008).

47. L. van der Maaten, “drtoolbox: Matlab Toolbox for DImensionality Reduction,” (2019).

48. P. Morel, “Gramm: Grammar of Graphics Plotting in Matlab,” The Journal of Open Source Software 3(23), 568 (2018).

49. D. Bates et al., “nlme: Linear and Nonlinear Mixed Effects Models,” (2018).

50. R. V. Lenth, “Least-Squares Means: The R Package lsmeans,” Journal of Statistical Software 69(1), 1–33 (2016).

51. R. D. Gibbons, D. Hedeker, and S. DuToit, “Advances in analysis of longitudinal data,” Annu Rev Clin Psychol 6(79–107 (2010).

52. X. A. Harrison et al., “A brief introduction to mixed effects modelling and multi-model inference in ecology,” PeerJ 6(e4794–e4794 (2018).

53. R. Hamid et al., “Comparison of alamar blue and MTT assays for high through-put screening,” Toxicol In Vitro 18(5), 703–710 (2004).

54. L. Lü et al., “Exocytosis of MTT formazan could exacerbate cell injury,” Toxicol In Vitro 26(4), 636–644 (2012).

55. G. Fotakis, and J. A. Timbrell, “In vitro cytotoxicity assays: comparison of LDH, neutral red, MTT and protein assay in hepatoma cell lines following exposure to cadmium chloride,” Toxicol Lett 160(2), 171–177 (2006).

56. A. van Tonder, A. M. Joubert, and A. D. Cromarty, “Limitations of the 3-(4,5-dimethylthiazol-2-yl)-2,5-diphenyl-2H-tetrazolium bromide (MTT) assay when compared to three commonly used cell enumeration assays,” BMC Res Notes 8(47 (2015).

57. S. H. Lee et al., “Tumor Evolution and Drug Response in Patient-Derived Organoid Models of Bladder Cancer,” Cell 173(2), 515–528 e517 (2018).

58. J. Xu, and W. Ying, “Increased green autofluorescence is a marker for non-invasive prediction of H2O2-induced cell death and decreases in the intracellular ATP of HaCaT cells,” bioRxiv 298075 (2018).

59. J. Surre et al., “Strong increase in the autofluorescence of cells signals struggle for survival,” Scientific Reports 8(1), 12088 (2018).

60. M. A. Borten et al., “Automated brightfield morphometry of 3D organoid populations by OrganoSeg,” Scientific Reports 8(1), 5319 (2018).

61. S. M. Tirier et al., “Pheno-seq – linking visual features and gene expression in 3D cell culture systems,” Scientific Reports 9(1), 12367 (2019).

62. G. Verbeke et al., “The analysis of multivariate longitudinal data: a review,” Stat Methods Med Res 23(1), 42–59 (2014).

63. S. M. H. Kashfi et al., “Morphological alterations of cultured human colorectal matched tumour and healthy organoids,” Oncotarget 9(12), 10572–10584 (2018).

