## Supplemental Material for "Patient-derived cancer organoid tracking with widefield one-photon redox imaging to assess treatment response"

Supplemental Table 1 – Definition of Quantitative Variables

| Variable Name | Variable Type | Definition (units) |
|---|---|---|
| **Organoid Area** | Morphological: Size | Number of pixels of segmented organoid multiplied by the squared pixel size ($mm^2$). |
| **Major Axis Length** | Morphological: Size | Length of major axis of ellipse that fits the segmented organoid (i.e., normalized second image moment) multiplied by the pixel size (mm). |
| **Minor Axis Length** | Morphological: Size | Length of minor axis of ellipse that fits the segmented organoid (i.e., normalized second image moment) multiplied by the pixel size (mm). |
| **Eccentricity** | Morphological: Shape | The ratio of the distance between the foci of the ellipse and its major axis length. An ellipse with eccentricity of 0 is a circle. An ellipse with eccentricity of 1 is a line. |
| **Circularity** | Morphological: Shape | $(4\pi * Organoid\ Area)/(Organoid\ Perimeter^2)$. For perfect circles, the value is 1. |
| **Convex Area** | Morphological: Size | Number of pixels in the convex hull containing the organoid multiplied by the squared pixel size ($mm^2$). |
| **Maximum Feret's Diameter** | Morphological: Size | Maximum distance between two antipodal vertices of the convex hull (i.e., caliper diameter) multiplied by the pixel size (mm). |
| **Minimum Feret's Diameter** | Morphological: Size | Minimum distance between two antipodal vertices of the convex hull (i.e., caliper diameter) multiplied by the pixel size (mm). |
| **Equivalent Diameter** | Morphological: Shape | Diameter of a circle with the same area as the organoid defined as $(4 * Area)/(\pi)$ multiplied by the pixel size (mm). |
| **Solidity** | Morphological: Shape | Proportion of the pixels in the convex hull that are also in the organoid defined as $(Area)/(Convex\ Area)$. |
| **Extent** | Morphological: Shape | Ratio of pixels in the organoid to pixels in the total bounding box defined as $(Area)/(Bounding\ Box\ Area)$. |
| **Organoid Perimeter** | Morphological: Size | Distance around the boundary of the organoid multiplied by the pixel size (mm). |
| **Optical Redox Ratio Mean** | Metabolic | Mean value of all redox ratio pixels in the organoid. |
| **Optical Redox Ratio Minimum** | Metabolic | Minimum value of all redox ratio pixels in the organoid. |
| **Optical Redox Ratio Maximum** | Metabolic | Maximum value of all redox ratio pixels in the organoid. |
| **Optical Redox Ratio Standard Deviation** | Metabolic | Standard deviation of all redox ratio pixels in the organoid. |
| **NAD(P)H Intensity Mean** | Metabolic | Mean value of all NAD(P)H pixels in the organoid. |
| **NAD(P)H Intensity Minimum** | Metabolic | Minimum value of all NAD(P)H pixels in the organoid. |
| **NAD(P)H Intensity Maximum** | Metabolic | Maximum value of all NAD(P)H pixels in the organoid. |
| **NAD(P)H Intensity Standard Deviation** | Metabolic | Standard deviation of all NAD(P)H pixels in the organoid. |
| **FAD Intensity Mean** | Metabolic | Mean value of all FAD pixels in the organoid. |
| **FAD Intensity Minimum** | Metabolic | Minimum value of all FAD pixels in the organoid. |
| **FAD Intensity Maximum** | Metabolic | Maximum value of all FAD pixels in the organoid. |
| **FAD Intensity Standard Deviation** | Metabolic | Standard deviation of all FAD pixels in the organoid. |

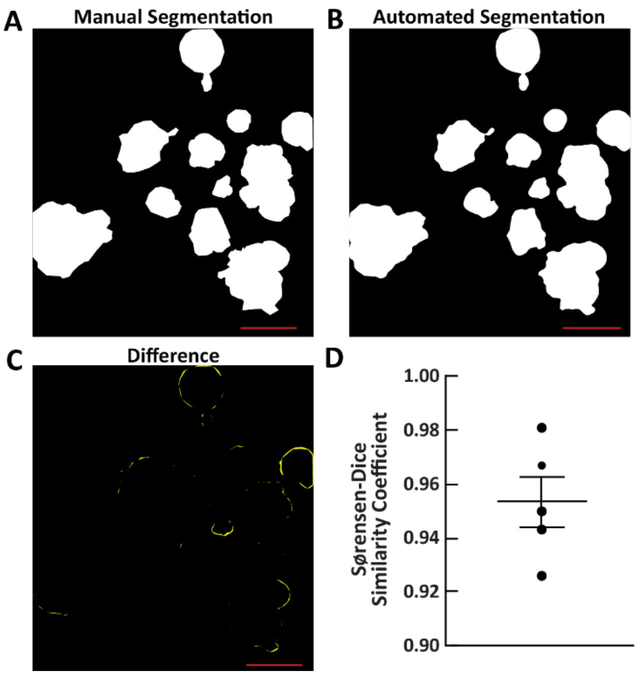


**Supplemental Figure 1. Validation of the Segmentation Pipeline.** The segmentation pipeline was validated by comparing the results of the automated pipeline to manual segmentation. (A) A representative image segmented manually in ImageJ. (B) The corresponding image after automated segmentation. (C) The difference between the automated and manual segmented images. (D) The Sørenson-Dice similarity coefficient for n = 5 image pairs plotted here as individual data points along with the mean and standard error. Scale bar: 500µm.

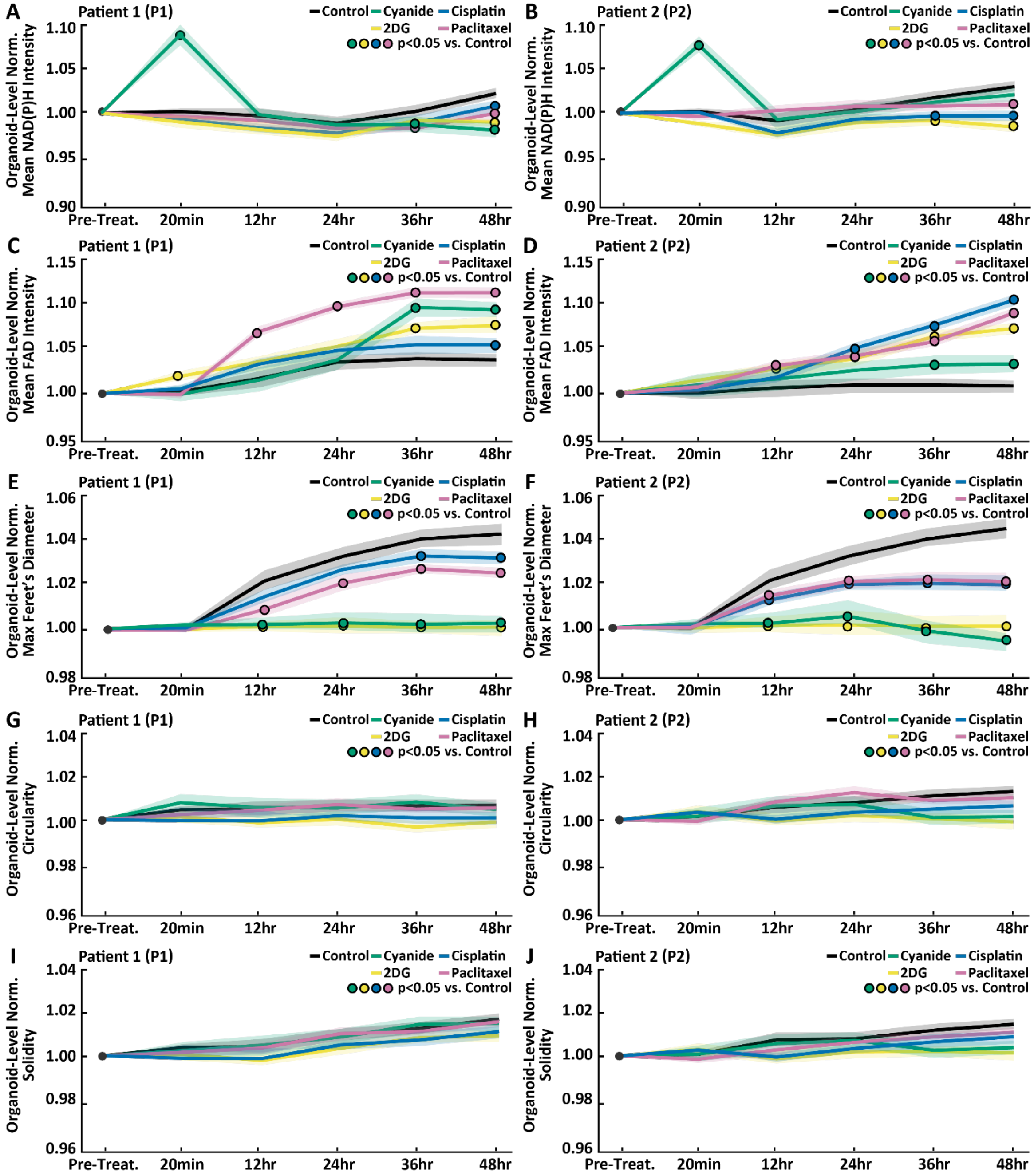


**Supplemental Figure 2. Time Series of Treatment-induced Changes in Additional Redox Imaging Variables.** A subset of the redox imaging variables is shown here. (A,B) Pre-treatment-normalized mean NAD(P)H intensity time series for all treatment groups for P1 and P2, respectively. (C,D) Pre-treatment-normalized mean FAD intensity time series for all treatment groups for P1 and P2, respectively. (E,F) Pre-treatment-normalized max Feret's diameter time series for all treatment groups for P1 and P2, respectively. (G,H) Pre-treatment-normalized circularity time series for all treatment groups for P1 and P2, respectively. (I,J) Pre-treatment-normalized solidity time series for all treatment groups for P1 and P2, respectively. Data plotted is mean ± standard error of the mean. Significant differences between a treatment and control is indicated with a colored circle for $p < 0.05$. Table 1 shows the number of organoids in each group and for each patient.

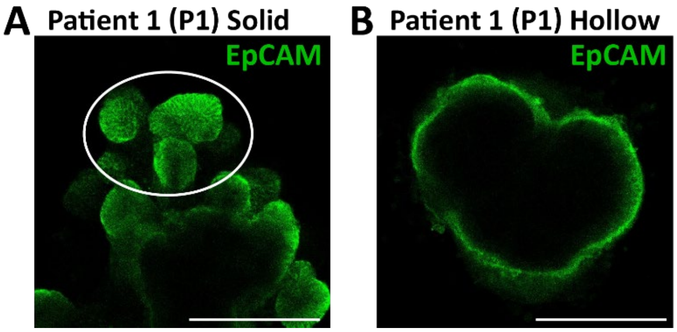


**Supplemental Figure 3. Confocal Microscopy of P1 Patient-Derived Cancer Organoids.** Confocal microscopy of patient-derived cancer organoids stained for epithelial cell adhesion molecular (EpCAM) illustrates the difference in the internal structure of P1 Solid and P1 Hollow patient-derived cancer organoids. (A) Confocal image of a P1 Solid patient-derived cancer organoid. The ellipse shows a region of densely organized cells characteristic of this phenotype. (B) Confocal image of a P1 Hollow patient-derived cancer organoid. Cells in these organoids are organized into a hollow shell. Scale bar: 100µm
